# Class-I myosin responds to changes in membrane tension during clathrin-mediated endocytosis in human induced pluripotent stem cells

**DOI:** 10.1101/2025.11.12.688091

**Authors:** Samantha L. Smith, Tong Zhan, Wan Li, Henry De Belly, Qing Zhang, Ke Xu, Orion D. Weiner, David G. Drubin

## Abstract

Clathrin-mediated endocytosis (CME) is an essential cellular process that needs to operate efficiently across a wide range of conditions. Internalization of the endocytic site involves forces generated by membrane-bound proteins and Arp2/3-mediated branched actin filament assembly to bend the plasma membrane (PM) from flat to omega-shaped. In mammalian CME, the requirement for a branched actin filament network varies depending on cell type and differences in membrane tension. However, how the actin network adapts to changes in load in order to ensure robustness of this process over a range of membrane tensions is not understood. Here, we combine live-cell imaging and super-resolution microscopy of genome-edited human induced pluripotent stem cells (hiPSCs) to investigate the role of the mammalian class-I myosin, Myosin1E (Myo1E), in load adaptation. Under normal conditions, sites that recruit Myo1E are rare and exhibit slow CME dynamics. However, as membrane tension increases and CME dynamics are slowed globally, Myo1E is recruited to more sites, likely to assemble more branched actin, resulting in increased force generation to rescue stalled sites and promote internalization. Loss of Myo1E results in increased Arp2/3 complex lifetime at CME sites under normal conditions, and at high membrane tension, these sites fail to recruit as many Arp2/3 molecules. We propose that Myo1E is recruited to CME sites that have stalled due to increased membrane tension, where it helps build a more effective branched actin network by generating force through motor activity and recruiting additional Arp2/3 complexes to rescue stalled sites.

**Significance:** For mammalian cells to internalize extracellular cargo via clathrin-mediated endocytosis (CME), specific regions of the plasma membrane (PM) must bend from flat to inwardly curved, a process that requires force-generating proteins. One key component in generating this force during CME is the branched actin network, in which actin filaments polymerize against the plasma membrane. When PM tension increases, more force is required to generate curvature, prompting the assembly of actin and actin associated proteins to aid the process. We demonstrate that the class-I myosin motor, Myosin1E (Myo1E), becomes increasingly crucial as membrane tension rises, presumably to build a more effective branched actin network to facilitate internalization of slowed sites.

## Introduction

Clathrin-mediated endocytosis (CME) is an evolutionarily conserved pathway through which cells internalize extracellular material, such as nutrients, signaling molecules, and transmembrane proteins, into vesicles that are trafficked throughout the cell^1^. Several key protein modules are responsible for enabling CME’s completion; the coat module, which captures cargo and bends the plasma membrane (PM) to form the vesicle, the actin module, which contributes to the force needed to create the invagination, and the scission module, which pinches off the formed vesicle from the plasma membrane for transport through the cytoplasm.

For cargo to be internalized through endocytosis, the plasma membrane must bend to make an invagination, which requires tens to thousands of pN, depending on the cell type^2,3^. The force to bend the plasma membrane is predominantly provided by membrane curvature-generating proteins and actin filament polymerization, which overcome forces from hydrostatic pressure and adhesion of the PM to the underlying cytoskeleton^4–7^. Many cellular processes involving membrane bending are assisted by the assembly of a branched actin filament network, nucleated by the Arp2/3 complex. Both *in vivo* and *in vitro* studies have demonstrated that the actin network adapts to high load through increased branching to provide greater force^8,9^.

The requirement for actin assembly at CME sites differs between yeast and mammals, as well as between different mammalian cell types and even different membrane regions of the same mammalian cell^10–12^. The first connection between actin-associated proteins and endocytic proteins was discovered using live cell imaging in mammalian cells and budding yeast^13–15^. The requirement for actin assembly at CME sites in yeast seems to be a consequence of the high turgor pressure, which is 0.2 – 1 MPa, as compared to ∼1 kPa in mammalian cells^10,16,17^. Mammalian cells do not experience turgor pressure, as they lack a cell wall, but experience osmotic and hydrostatic pressure^18,19^. The largest energetic barrier that mammalian cells must overcome to internalize plasma membrane during CME is membrane tension^2^. When mammalian cells are treated with actin-perturbing drugs, the effects on CME are highly variable, which initially made demonstrating a clear function for actin in CME challenging. Several lines of evidence have now established that dependence on actin assembly during CME increases as membrane tension increases in mammalian cells^5,7,10^. When mammalian cells were placed under hypotonic shock or stretched, resulting in increased membrane tension, and treated with actin-perturbing drugs, CME proteins had increased lifetimes, and internalization was stalled compared to when no perturbing drugs were added^5^. Additionally, in mammalian cells under hypotonic shock, more actin covered the clathrin-coated surface compared to isotonic conditions^7^. Recent evidence also demonstrated that the Arp2/3 complex is positioned to stimulate asymmetric actin assembly and force production^20^. Actin’s response to increased membrane tension has proven to be pivotal to robust CME under varied membrane tensions; what triggers this response is less clear.

One mechanism cells use to respond to varying membrane tension is changing membrane-to-actin cortex attachments^21^. In CME, a potential candidate that could fulfill this role is a class-I myosin motor. Class-I myosins are single-headed, actin-based motors involved in membrane dynamics and trafficking^22^. The involvement of class-I myosin motors in endocytosis was first discovered in yeast, where the motors are termed Myo3 and Myo5. Deletion of the genes encoding these proteins caused substantial defects in CME^23,24^. More specifically, point mutations in either the motor domain that binds actin filaments or mutations in the Arp2/3 nucleation domain of Myo5 resulted in defective, stalled endocytic sites^25^. Subsequently, the mammalian homolog of Myo3 and Myo5, Myosin1E (Myo1E), was found to participate in mammalian endocytosis, where it binds to synaptojanin and dynamin2, two endocytic proteins that drive membrane scission^26^. Myo1E is a member of the unconventional class-I myosin family. Myo1E consists of multiple protein domains, most importantly, an N-terminal ATPase motor domain that binds actin filaments, a TH1 domain that binds to anionic phospholipids, and an SH3 domain that binds to actin nucleation-promoting factors (NPFs), dynamin2 (Dnm2), and synaptojanin^26–28^. Since this myosin can bind to both actin filaments and the PM simultaneously, it has the capacity to act as a bridge between the two, potentially modulating membrane-to-actin cortex attachment adaptation and facilitating actin branching through its SH3 domain.

Here, we tested the hypothesis that Myo1E participates in load adaptation in response to changes in membrane tension by live-cell imaging of triple-genome-edited human induced pluripotent stem cells (hiPSCs). Endocytic dynamics were quantified under normal physiological conditions and under increased membrane tension to investigate Myo1E’s role in CME. Endocytic events were analyzed in an unbiased manner using single particle tracking algorithms to dissect differences between protein dynamics within the control stem cells, cells with elevated membrane tension, and cells with the gene for Myo1E knocked out. Our results demonstrate that, under normal conditions, Myo1E’s recruitment to CME sites is rare in hiPSCs. However, as membrane tension increases, the fraction of endocytic sites that recruit Myo1E dramatically increases. Interestingly, under normal conditions, sites that recruit Myo1E appear to be longer-lived, which may hint at these sites having unique local environmental signatures that create a requirement for Myo1E to aid in internalization. Additionally, as tension increases, sites that do and do not recruit Myo1E behave more similarly and seem more stable, moving less within the plasma membrane plane and with longer endocytic protein lifetimes.

## Results

### Myo1E is a late-arriving CME protein that localizes asymmetrically at endocytic sites in hiPSCs

To analyze the spatial dynamics of Myo1E during mammalian CME, we first endogenously tagged MYO1E with a HaloTag using CRISPR/Cas9 genome editing in hiPSCs, in a cell line previously genome-edited to express two tagged endocytic proteins: AP2-tagRFP-T, a canonical endocytic coat marker, and DNM2-tagGFP2, the scission factor (tagged proteins referred to as AP2-RFP, Dnm2-GFP, and Myo1E-JF635) (Fig. 1A, S1A, Movie 1). After genome editing, triple-tagged clones were screened for lines with endocytic dynamics profiles matching those of the parental cell line (Fig. S1B-G). The appearance of AP2-RFP as a punctum on the plasma membrane marks the start of an endocytic event, while the peak of Dnm2-GFP marks vesicle scission^20^. These markers allowed us to determine when Myo1E is recruited relative to the initiation and completion of endocytosis. We confirmed previous reports indicating that Myo1E is a late-arriving endocytic protein that appears in a short burst near the end of CME events, with a lifetime of approximately 7.6 seconds (Fig. 1B)^26,27^.

**Figure 1.**
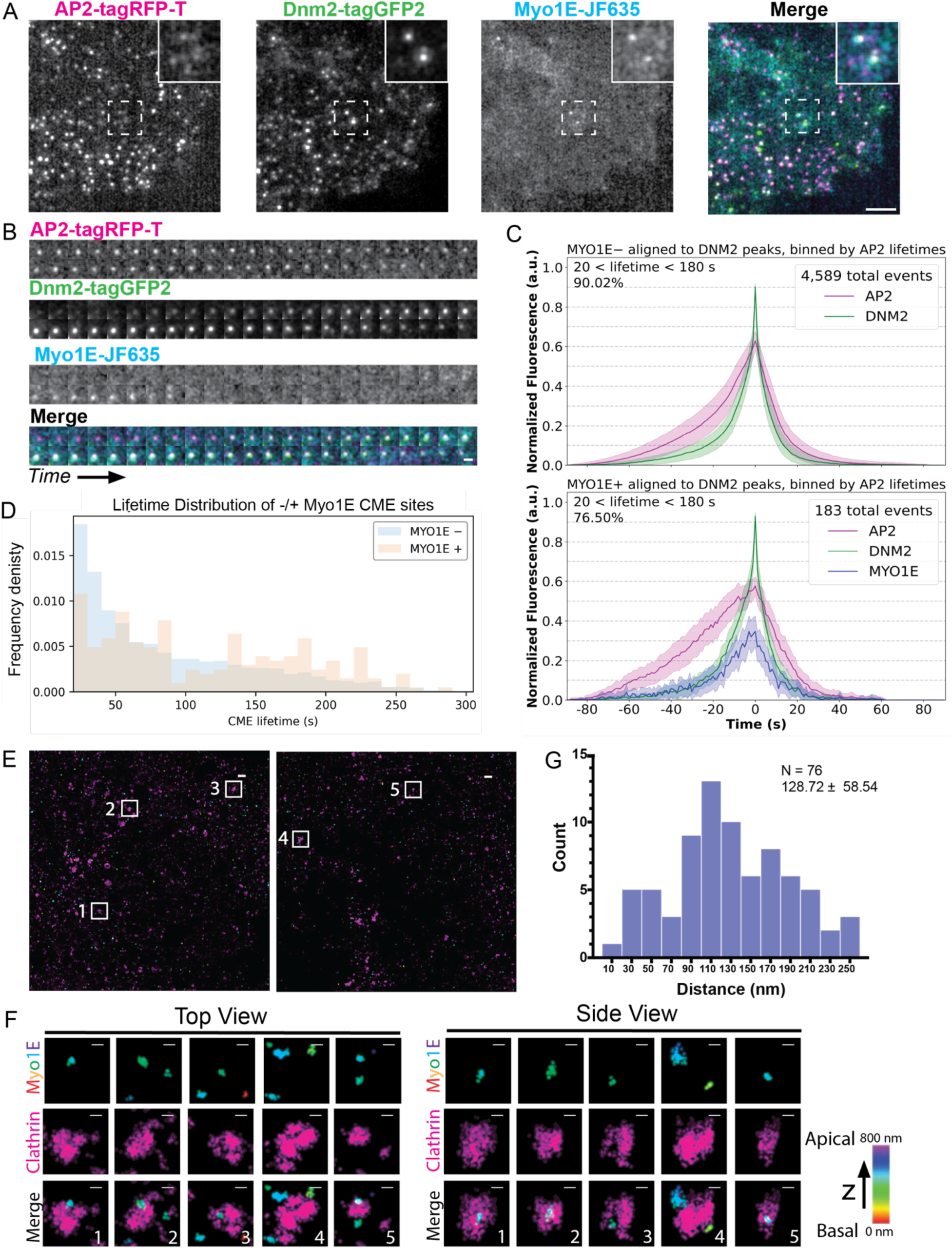
(A) Total Internal Reflection Fluorescence (TIRF) still image from a 5-minute movie imaged at 1 frame per second of genome-edited AP2-tagRFP-T; DNM2-tagGFP2; Myo1E-JF635 (ADM) hiPSCs. Scale bar = 5 µm. (B) Montage of an endocytic event where each image is one second from a 5-minute movie. The mean lifetime of Myo1E = ∼7.6 seconds (n= 563 sites). Scale bar = 1 µm. (C) Averaged intensity versus time plots of Myo1E-negative and Myo1E-positive CME site within the ADM cell line. These traces represent the fluorescent signal of each protein for a single endocytic event averaged across n-events. The fluorescent intensity is normalized to Dnm2 intensity. The lifetimes of the events plotted are between 20 and 180 seconds for 90.02% of the Myo1E-negative sites and 76.50% of the Myo1E-positive sites. Error bar: ¼ standard deviation. Events are aligned to the frames showing the maximum Dnm2-GFP intensity (time = 0 sec). (D) Frequency distribution plot of Myo1E-negative (n = 6,612) and Myo1E-positive sites (n = 560).This distribution highlights that Myo1E-positive events tend to have longer endocytic lifetimes than sites that do not recruit Myo1E. (E) Two-color 3D STORM image of the bottom membrane of ADM cells immunolabeled with clathrin heavy chain antibody (magenta) and HaloTag (rainbow, correlating to depth in z). Scale bar = 500 nm. (F) Zoom in image of the numbered white boxes in (E). Scale bar = 1 00 nm. (G) Histogram plot of the measured 2D distance between the centroids of the clathrin and Myo1E signal. The mean and standard deviation are reported on the graph (n = 76 sites).

Using a custom MATLAB single-particle tracking program, we analyzed 56,160 endocytic events, tracing how each protein was recruited over time^4,20,29^. We then applied strict criteria to identify true endocytic events, defined as those with lifetimes between 20 and 300 seconds. Of the CME sites with lifetimes between 20 and 300 seconds, only ∼8.7% recruited detectable Myo1E, indicating that Myo1E recruitment to CME sites is quite rare in hiPSCs. Ninety percent of the CME sites that did not recruit detectable Myo1E had lifetimes between 20 and 180 seconds (n = 4,589 sites meeting this criterion) (Fig. 1 C, top). Of those CME sites that did recruit Myo1E, 76.5% of these had lifetimes between 20 and 180 seconds (n = 183 sites), with Myo1E recruitment slightly lagging behind Dnm2-GFP recruitment, and its disappearance preceding that of Dnm2-GFP (Fig. 1 C, bottom). The remaining 23.5% of Myo1E-positive sites had lifetimes between 180 and 300 seconds. Distribution analysis revealed that CME sites to which Myo1E was recruited tended to have longer lifetimes compared to those that do not (lifetimes without Myo1E = 84.5 ± 61. 3 seconds; with Myo1E = 114.5 ± 69.8 seconds) (Fig. 1 D). Myo1E was recruited to 17.3% of long-lived or persistent events, classified as events with lifetimes exceeding 300 seconds, substantially more than the 8.7% observed for non-persistent sites.

Seeing that Myo1E is preferentially recruited to longer-lived endocytic sites, we next asked where Myo1E localizes at CME sites. Since previous work showed that the Arp2/3 complex and its NPF, N-WASP, are located asymmetrically at CME sites, we investigated whether Myo1E has a similar localization pattern^20^. To address this we used 3D Stochastic Optical Reconstruction Microscopy (STORM) imaging on fixed AP2-RFP;DNM2-GFP;MYO1E-HaloTag hiPSCs, staining for clathrin and HaloTag^30,31^. Using two-color 3D STORM, we observed that Myo1E puncta also localize asymmetrically at CME sites (Fig.1E & F)^32^. We determined that Myo1E is about 128.72 ± 58.5 nm from the center of the clathrin coat (n = 76 sites) (Fig.1G). Together, these results confirm that Myo1E is a late-arriving protein in mammalian CME, whose recruitment shadows the arrival of Dnm2, but additionally show that Myo1E tends to be recruited to longer-lived endocytic sites^27^. Additionally, Myo1E localizes asymmetrically at CME sites, similar to what has been observed for other proteins in the actin module (Arp2/3 and N-WASP)^20^.

### Loss of Myo1E alters Arp2/3 complex lifetime and its distribution at endocytic sites in hiPSCs

We next investigated how loss of Myo1E affects endocytosis. Using CRISPR/Cas-9, we generated a Myo1E homozygous knockout cell line to test for effects on endocytic dynamics (Fig.2A, S2A). Following gene editing, clones were screened to assay for defects in endocytic profiles compared to unedited Myo1E cell lines (Fig.S2B-E). We also tagged a single allele of the ARPC3 gene, encoding the ArpC3 subunit of the Arp2/3 complex, with HaloTag (hereafter called ArpC3-JF635) to determine how the branched actin filament module at CME sites responds to loss of Myo1E. Using the same particle tracking and analysis software mentioned above, we found that loss of Myo1E did not grossly affect CME dynamics or internalization of transferrin, a canonical CME cargo (Fig.2B, S2F, Movie 2 and 3). We next compared the number of active tracks, those with lifetimes between 20 and 300 seconds, and persistent tracks, those with lifetimes longer than 301 seconds. Notably, there were fewer active ArpC3-positive CME tracks in the Myo1E-/- knockout cell line compared to control cells (control = 75.1% ± 6.3, Myo1E-/- = 65.8% ± 14.0) (Fig.2C, S2G). Additionally, the average lifetime of ArpC3-JF635 increased from ∼25.5 seconds in control cells to ∼28.0 seconds in Myo1E knockout cells, consistent with the possibility that Myo1E is important in nucleating branched actin or in global membrane tension maintenance and/or in adaptation to local differences in membrane tension (Fig.2D)^33^. We next determined if there were any differences in the spatial displacement of endocytic sites from their origins as CME progresses in the presence or absence of Myo1E. In the AP2-RFP; Dnm2-GFP; Myo1E-JF635 cell line, we observed that sites that recruit Myo1E (hereafter Myo1E-positive) show a lower AP2-RFP mean squared displacement (MSD) (∼50.0 ± 25.6 nm^2^) than sites that do not recruit Myo1E (∼73.9 ± 32.5 nm^2^) (hereafter Myo1E-negative) (Fig.2E). However, in Myo1E-/- cells, the MSD of AP2-RFP, at sites that recruit ArpC3, hardly decreased compared to control cells (Myo1E-/- = ∼66.6± 29.3 nm^2^, control =∼68.3± 28.3 nm^2^), which may reflect that fact that most CME sites do not recruit Myo1E (Fig.2E). If we compare the AP2-RFP MSD at CME sites that recruit Myo1E in Myo1E+/+ cells (∼50.0 ± 25.6 nm^2^) to the MSD of AP2-RFP in Myo1E+/+ cells that recruit ArpC3 (∼68.3± 28.3 nm^2^), we also see a statistically significant difference (P <0.0001 for Mann-Whitney test), which means that sites that recruit Myo1E are less motile than sites that recruit branched actin.

**Figure 2.**
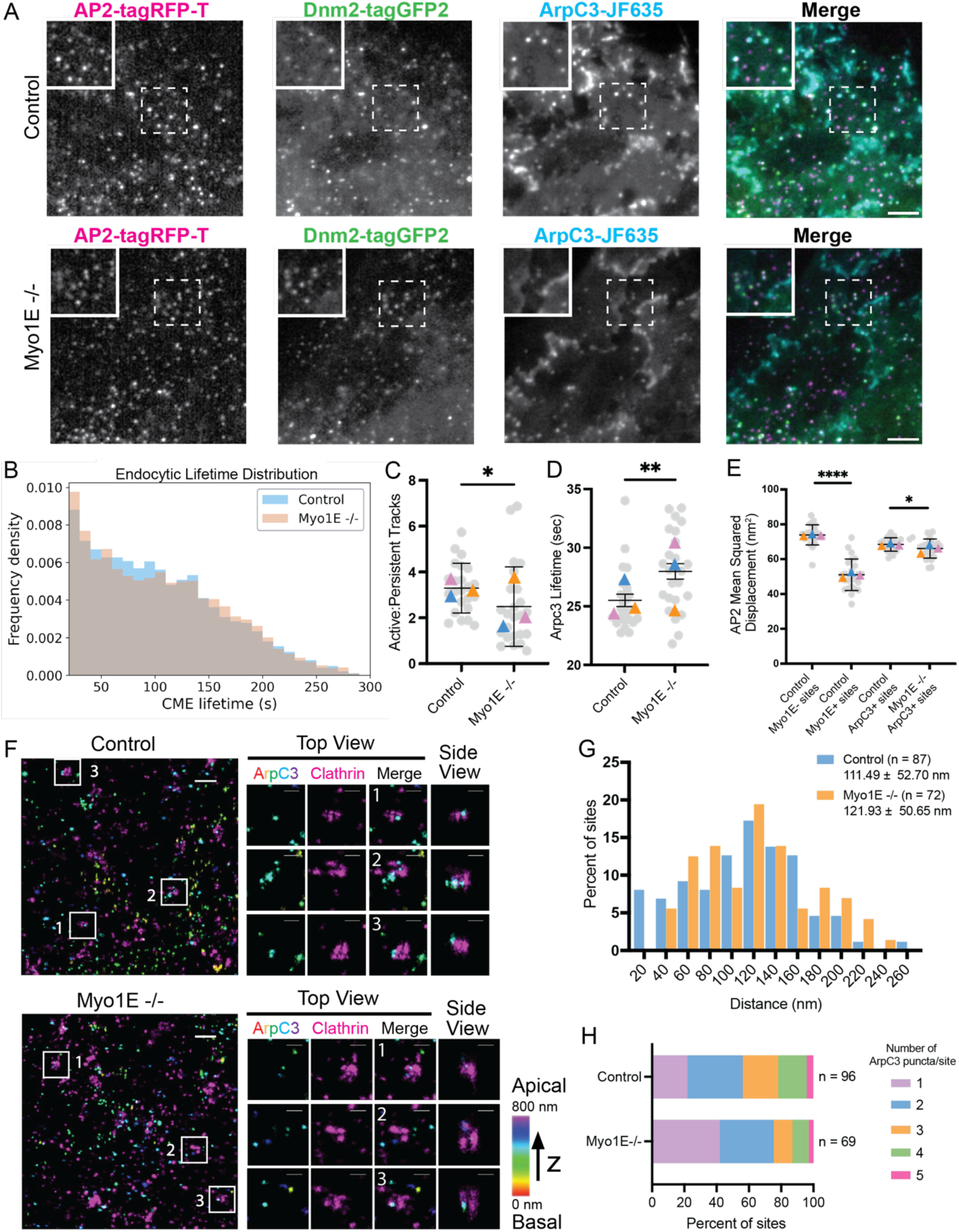
(A) TIRF still image of genome-edited AP2-tagRFP-T; DNM2-tagGFP2; ArpC3-JF635 (ADA) hiPSCs. TIRF image of genome-edited AP2-tagRFP-T; DNM2-tagGFP2; ArpC3-JF635; Myo1E knockout (ADAM) hiPSCs. Scale bar = 5 µm. (B) Cumulative frequency distribution plot of endocytic lifetimes in ADA cells (n = 5,911 sites) compared to ADAM cells (n = 5,608 sites). (C-D) Scatter plots where the grey circles represent the mean of each movie and the colored triangles are the mean of each technical replicate. The horizontal black line represents the mean, with the error bars showing the SD. (C) Scatter plot showing the ratio of active to persistent tracks between control ADA cells (n = 24 movies) and ADAM cells (n = 25 movies). Persistent tracks are classified as events that last longer than 301 seconds. (P = 0.0079, for the Mann-Whitney test). (D) Scatter plot showing the lifetime of ArpC3 at CME sites is increased in ADAM cells compared to control ADA cells. (P = 0.0022, for Mann-Whitney test). (E) Scatter plot showing the mean displacement of AP2-RFP. In control sites that recruit Myo1E in ADM cells, the mean displacement of AP2-RFP is reduced compared to sites that do not recruit Myo1E in ADM cells (P**** <0.0001, for the Mann-Whitney test). Endocytic sites that recruit ArpC3 in Myo1E knockout cells show a slight reduction in AP2-RFP mean displacement compared to ArpC3-positive sites in ADA cells (P = 0.0454, for the Mann-Whitney test). (F) Two-color 3D STORM image of the bottom membrane of control ADA cells immunolabeled with clathrin heavy chain antibody (magenta) and HaloTag (rainbow, correlating to depth in z). Scale bar = 500 nm. Zoomed-in image of the numbered white boxes (right). Scale bar = 200 nm. (G) Histogram of the measured distance between the centroids of the clathrin and ArpC3 signal. The distance between ArpC3 and clathrin increases in ADAM cells (P = 0.206, for t-Test with Welch Correction). (H) Stacked bar plot showing the number of ArpC3 puncta per clathrin site in control and ADAM cells.

Since we observed a difference in ArpC3 lifetimes in Myo1E-/- cells compared to control cells, we next examined whether branched actin filament network structure is affected by Myo1E knockout. Using the same 2-color 3D STORM approach as above, we observed that the Arp2/3 complex is still asymmetric on the clathrin coat even in the absence of Myo1E (Fig.2F). Interestingly, when measuring the distance from the center of the clathrin coat to the center of the ArpC3 puncta, we observed an increase in this distance in the Myo1E knockout cells (control = 111.5 ± 52.7 nm (n = 87), Myo1E-/- = 121.9 ± 50.7 nm (n = 72)) (Fig.2G). We also found that, in Myo1E-/- cells, fewer ArpC3 puncta were recruited to clathrin-coated pits (CCPs) (Fig.2H). We hypothesize that, under normal conditions, Myo1E-/- cells may build less efficient branched actin filament networks, explaining the increased ArpC3 lifetimes and increased distance between ArpC3 puncta and clathrin, but still manage to complete endocytosis.

### Increased membrane tension, as a result of micro-aspiration, leads to dramatic re-localization of Myosin1E and the Arp2/3 complex

While loss of Myo1E does not appear to drastically alter endocytic dynamics, but does appear to affect branched actin filament dynamics, we wondered if Myo1E recruitment to only a subset of CME sites might be the result of differences in local membrane tension at CME sites. Previous reports demonstrated that the branched actin filament network responds to changes in membrane tension by increasing actin assembly^5,7^. To assess how Myo1E responds to a sustained increase in membrane tension, we used micro-aspiration to pull on the cell membrane and mechanically raise membrane tension^34,35^. During live-cell imaging, membrane tension was raised by attaching a micro-aspirating pipette to the edge of a single cell within a stem cell colony and aspirating until the membrane appeared inside the pipette. To assess the effect of pulling, we first determined how the Arp2/3 complex responds to this method of mechanically increasing membrane tension. We observed a sharp increase in ArpC3 signal at the cell boundaries as well as at CME sites after aspirating the cells (Fig.3A, bottom, Movie 4). After aspiration, the ArpC3 fluorescent signal across the whole cell was, on average, 9.7% higher than prior to aspiration, and in some cells increased by as much as 30% (Fig.3B and C). This well-established method of increasing the membrane tension has previously been shown to have profound effects on the cortical branched actin network and actin-associated proteins^36,37^. As previously reported, we observe that the fluorescent ArpC3 signal increases and then plateaus, remaining above the initial fluorescent signal prior to pulling (Fig.3B).

**Figure 3.**
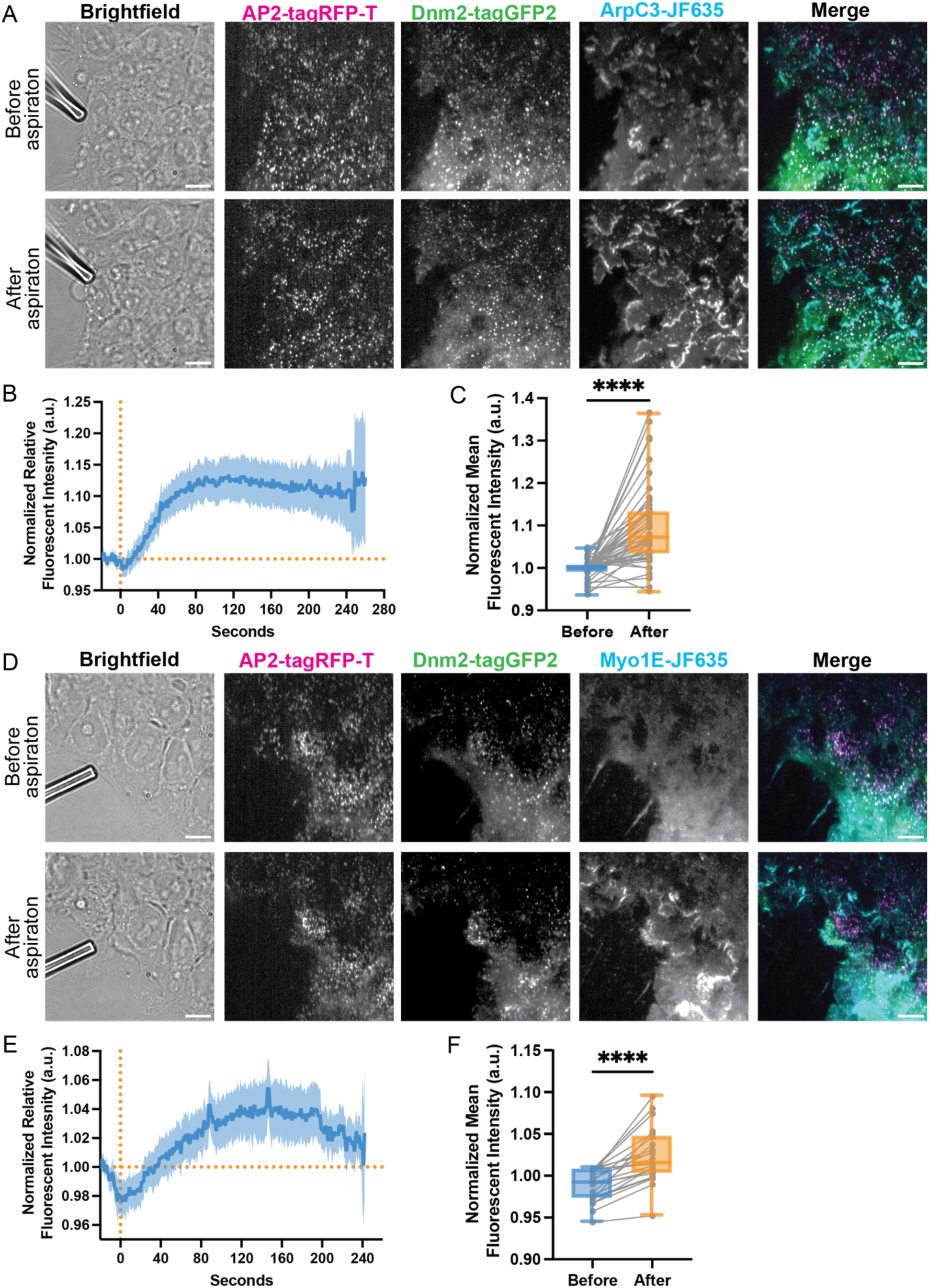
(A) TIRF image of ADA cells before and after aspiration Scale bar = 10 µm. (B) Trace of normalized ArpC3 fluorescent intensity over time of each cell the cell being pulled on, where the solid blue line represents the mean and the light blue represents the error with 95% CI (n = 51 cells). (C) Box and whisker plot of the mean fluorescent intensity of ArpC3 for the time before and the time after aspiration (P**** < 0.0001, for the Wilcoxon test). (D) TIRF image of ADM cells before and after aspiration. Scale bar = 10 µm. (E) Trace of normalized Myo1E fluorescent intensity over time of each cell, where the solid blue line represents the mean and the light blue represents the error with 95% CI (n = 23 cells). The vertical dashed orange line represents where aspiration started, and the horizontal line represents the baseline fluorescent intensity before aspiration, used for normalization. (F) Box and whisker plot of the mean fluorescent intensity of Myo1E for the time before and the time after aspiration. The grey dots represent the individual mean values for each cell connected with the mean values after aspiration for the same cell (P**** < .0001, for the Wilcoxon test).

We next tested whether Myo1E undergoes changes membrane localization in response to increased membrane tension. Before aspiration, Myo1E was largely diffuse across the plasma membrane with few puncta, some of which associated with CME sites (Fig.3D, top, Movie 5). After aspiration, Myo1E dramatically redistributed to the cell boundaries, and more puncta appeared at both CME and non-CME punctate sites (Fig.3D, bottom, Movie 5). Similar to ArpC3 signal over time, Myo1E signal increased, on average about 4.8% across the entire cell, after aspiration and plateaued (Fig.3E). The mean fluorescence intensity of Myo1E was higher per cell after aspiration compared to the fluorescence signal before aspiration (Fig.3F). To our knowledge, the results from this assay are the first to establish that Myo1E responds to changes in membrane tension and does so by redistributing to cell boundaries where membrane tension is likely highest as the neighboring cells are pulled away from one another. These results indicate that Myo1E responds to changes in membrane tension by being recruited to sites of enriched in branched actin filaments.

### Hypotonic shock increases the proportion of endocytic sites that recruit Myosin1E

Micro-aspiration allowed us to visualize the global response of ArpC3 and Myo1E to increased membrane tension through mechanical manipulation. To have more control over a range of membrane tensions, we turned to hypotonic shock. Cells were swelled to varying degrees by decreasing the osmolarity of the imaging solution to concentrations of 300 mOsm (no shock), 225 mOsm, 150 mOsm, and 75 mOsm (high shock). By live-cell imaging, we observed a dramatic increase in Myo1E localization to endocytic sites upon osmotic shock induction (Fig.4A, S3A, Movie 6 and 7). In alignment with our micro-aspiration assay, prior to shock we observed Myo1E diffuse across the plasma membrane, and upon shock induction Myo1E redistributed to cell boundaries, but also more puncta appeared. As membrane tension increased, there was an increase in endocytic lifetime from ∼151.5 seconds to ∼195.5 seconds in response to 75 mOsm media, the highest shock condition (Fig.S3B). At the highest membrane tension state, 75 mOsm, there was also an increase in the number of persistent sites compared to the other states (Fig.4B). We observe a dose-dependent response in the number of Myo1E positive puncta relative to tension, with control cells having Myo1E at approximately 8% of sites, rising to about 33% at 75 mOsm (Fig.4C). We quantified CME site initiation rates by measuring the total number of sites formed during a 5-minute movie across the area of the cell colony. Our results reveal a 2.8-fold increase in the initiation of Myo1E-positive sites as membrane tension increases from 0.0007 ± 0.0004 events/µm^2^/min in control cells to 0.002 ± 0.002 events/µm^2^/min at 75 mOsm (Fig.4D). These results demonstrate that Myo1E recruitment to endocytic sites as well as the likelihood that a newly formed site will have Myo1E increases as membrane tension increases.

**Figure 4.**
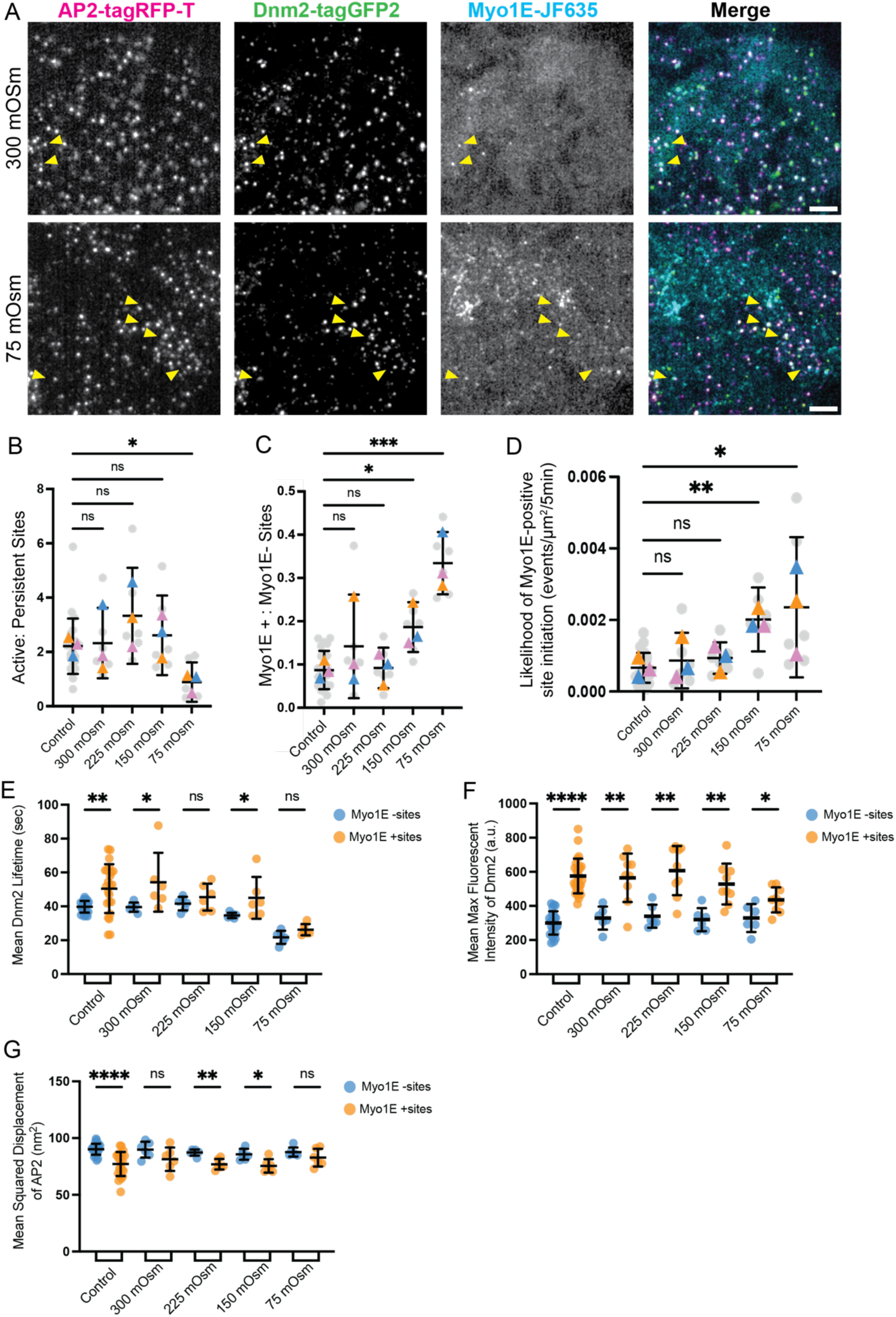
(A) TIRF image of ADM cells after the addition of 300 mOsm media and after the addition of hypotonic media solution to reach a final concentration of 75 mOsm. Yellow arrowheads indicate examples in which AP2-RFP, Dnm2-GFP, and Myo1E co-localize. Scale bar = 5 µm. (B) Scatterplot showing the ratio of active to persistent Myo1E-positive sites. (P = 0.0165, for Kruskal-Wallis test). (C) Scatterplot showing the ratio of Myo1E-positive sites to Myo1E-negative sites per movie, in grey dots, and the mean of each replicate, in colored triangles. Control conditions are those for which the media was not changed, and 300 mOsm conditions are those where media with the same osmolarity as the imaging media was added. (P* = 0.0215, P*** = 0.0001, for Kruskal-Wallis test). (D) Scatterplot showing the likelihood of site initiation (initiation rate) for Myo1E-positive sites. (P** = 0.0063, P* = 0.0226, for Kruskal-Wallis test). (E-G) Scatterplots characterizing (E) Dnm2-GFP lifetime, (F) mean fluorescent intensity, and (G) mean displacement between endocytic sites that are Myo1E negative or Myo1E positive (P**** < 0.0001, for Mann-Whitney). All error bars show SD. For control conditions, n = 24 movies, and for osmotic shock conditions, n = 6 movies, for all quantifications.

We also determined whether Dnm2-GFP, a direct binding partner of Myo1E, undergoes changes in response to increased membrane tension^26^. In control conditions and at 300 mOsm, the average lifetime of Dnm2-GFP was ∼39.7 seconds for Myo1E-negative sites and ∼50.9 seconds for Myo1E-positive (Fig. 4E). Similarly, the average fluorescence intensity of Dnm2-GFP was almost twice as high at Myo1E-positive sites (575.8 ± 107.1 a.u.) as at Myo1E-negative sites (300.5 ± 68.3 a.u.) (Fig.4F). The increase in Dnm2-GFP fluorescence intensity at Myo1E-positive sites suggests that almost twice as many Dnm2 subunits are present at these sites, consistent with the possibility that additional scission force is required to pinch off vesicles at these sites. As previously mentioned, we observed that the MSD of AP2 decreased from 90.2 nm^2^ at Myo1E-negative sites to 77.2 nm^2^ at Myo1E-positive sites, under normal growth conditions, highlighting the increase in vesicle stability in the presence of Myo1E (Fig.2E & 4G). At 75 mOsm, the differences in the Dnm2-GFP lifetime and fluorescence intensity, as well as the MSD of AP2 in Myo1E-negative and Myo1E-positive sites, showed little to no difference (Fig.4E-G). Also, we observed that the AP2-RFP lifetime was higher across all osmotic shock conditions in the Myo1E-positive sites compared to Myo1E-negative sites (Fig.S3C). Interestingly, the amount of AP2-RFP recruited to both Myo1E-negative and Myo1E-positive sites was similar, indicating that recruitment of CME coat proteins does not change in response to hypotonic shock (Fig.S3D). In contrast to the AP2-RFP and Dnm2-GFP MSD showed little to no difference between Myo1E-negative and Myo1E-positive sites, possibly because Dnm2-GFP is associated with the plasma membrane rather than the invaginating membrane and is under finer constraint until scission is completed (Fig.S3E). These findings suggest that under normal conditions, endocytic sites to which Myo1E has been recruited are distinct from those that do not and may represent stalled sites that require additional endocytic proteins to proceed to scission. This distinction between Myo1E-positive and Myo1E-negative sites is diminished when membrane tension is elevated because more sites are stalled and recruit Myo1E.

### In cells lacking Myo1E, fewer Arp2/3 complex molecules are recruited to CME sites in response to increased membrane tension, and CME sites show greater lateral motility

Our osmotic shock assay showed that, regardless of membrane tension, Myo1E-positive CME sites have increased endocytic lifetimes and recruit more endocytic proteins. Additionally, as membrane tension increases, Myo1E-positive and negative sites behave more similarly than under isotonic conditions. We next investigated how loss of Myo1E influences Arp2/3 complex recruitment and dynamics over a range of membrane tensions. As with our micro-aspiration results, the ArpC3 signal at CME sites in control cells rose sharply upon shock induction, and ArpC3 was recruited to more CME sites under increased membrane tension, with 242±32 sites with ArpC3 per 5 min movie at 300 mOsm and 458 ± 130 sites with ArpC3 per 5 min movie at 75 mOsm (Fig.5A, Movie 8 and 9). In Myo1E knockout cells, ArpC3 was still recruited to cell edges and puncta in response to osmotic shock (Fig.5B, Movie 10 and 11). However, while there are more ArpC3 puncta, the puncta that are associated with CME sites dropped from 246 ± 22 sites per 5 min movie at 300 mOsm to 182 ± 43 sites per 5 min movie at 75 mOsm. When we measured the lifetime of ArpC3 at CME sites across various osmotic conditions, we found that the average lifetime of ArpC3 is longer in Myo1E knockout cells compared to controls (control = ∼33.4 seconds, Myo1E-/- = ∼38.7 seconds) (Fig.5C). Intriguingly, if we quantify the normalized mean fluorescence intensity of ArpC3 at active and persistent sites, the ArpC3 signal remained the same between control and knockout cells under all osmotic shocks, except at the highest osmotic shock level (control = ∼1.5 a.u., Myo1E-/- = ∼1.1 a.u.) (Fig.5D). Furthermore, the MSD of ArpC3 at active sites is higher in Myo1E knockout cells than in controls across all conditions (Fig.5E). These results suggest that, in hiPSCs lacking Myo1E, fewer CME sites assemble branched actin filaments, and that when branched actin filaments are nucleated at CME sites, less force is produced because fewer ArpC3 molecules are recruited under high osmotic shock.

**Figure 5.**
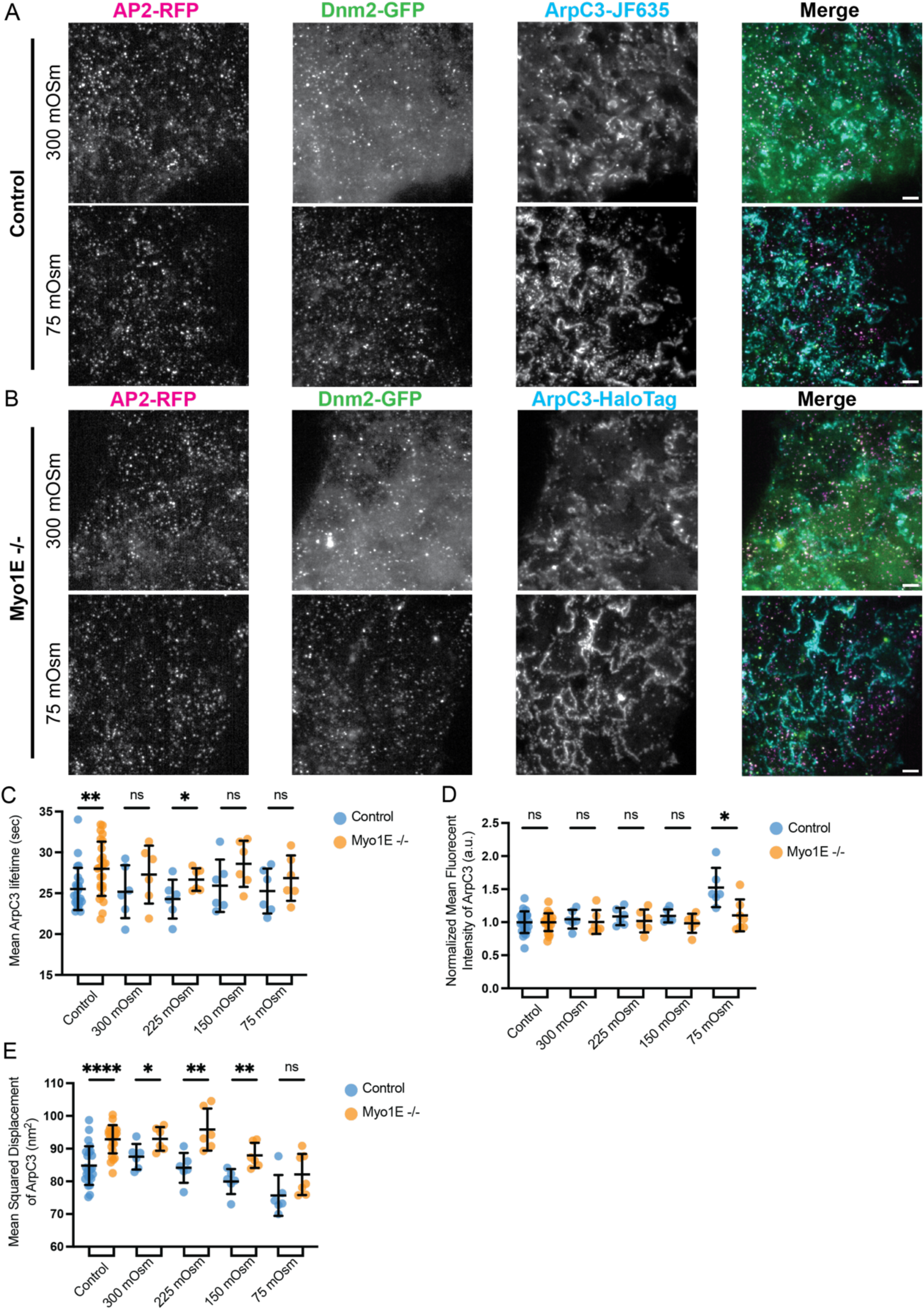
(A-B) TIRF image of ADA or ADAM cells after the addition of 300 mOsm media and after the addition of hypotonic media solution to reach a final concentration of 75 mOsm. Scale bar = 5 µm. (C-E) Scatterplots of (C) ArpC3 lifetime, (D) normalized mean fluorescent intensity, and (E) mean displacement from the center of endocytic sites in control cells and Myo1E knockout cells. Error bars show SD. (P**** < 0.0001, for Mann-Whitney) For control conditions, n = 24 movies and for shock conditions n = 6 movies for all quantifications.

## Discussion

While the role of type-1 myosin has been extensively investigated in yeast CME, the role of its ortholog, Myo1E, in mammalian CME has been less well studied. Here, we set out to investigate the function of Myo1E in hiPSCs. Previous work demonstrated that the dependency on actin filaments during CME in mammalian cells varies with cell type, plating substrate, cargo size, and membrane tension^4,5,7,38–40^. In addition, branched actin filaments are not assembled at every endocytic site, even within the same cell^20^. Against this background of heterogeneity across cell types and between CME sites within the same cell, we set out to investigate Myo1E’s role in CME.

Through genome editing of the MYO1E gene in hiPSCs previously engineered to express two fluorescent protein-tagged CME proteins (AP2 and DNM2) at endogenous levels, we were able to analyze thousands of CME events without bias. Consistent with previous findings, in hiPSCs, Myo1E is a late-arriving endocytic protein that is recruited to CME sites around the same time as Dnm2^27^. Interestingly, as observed previously for branched actin filaments, Myo1E is preferentially recruited to longer-lived CME sites, which is not surprising considering that Myo1E is part of the actin module and possesses an SH3 domain that recruits Arp 2/3 NPFs to CME sites ^20,27^. We speculate that the local environments at certain CME sites make CME completion more challenging. Myo1E may aid in completing CME through its SH3 domain by recruiting more NPFs to boost Arp2/3 activity, or through its motor domain to generate force for internalization, or both. Because Myo1E’s SH3 domain can recruit NPFs, which activate the Arp2/3 complex, we wondered if Myo1E is located asymmetrically at CME sites, like the Arp2/3 complex^20^. Earlier studies used average protein distribution and radial density profiles from STORM images to examine Myo5 localization at CME sites in yeast, concluding that Myo5 forms a continuous ring around them^41^. In our study, combining STORM with TIRF microscopy, we examined Myo1E localization across hundreds of sites and discovered that it exhibits a similar asymmetric localization pattern as the Arp2/3 complex in mammalian cells.

Unlike yeast, where type 1 myosin is essential for CME, knockout of the gene encoding Myo1E in iPSCs did not cause a gross defect in CME dynamics. However, Myo1E’s loss resulted in an increase in Arp2/3 complex lifetime at CME sites. A possible explanation for this observation is that the loss of this membrane-to-actin-cortex attachment across the entire cell perturbs force production by the branched actin network at CME sites. Thus, the branched actin filament network that assembles at CME sites must work longer to complete endocytosis. Our data also supports the possibility that the branched actin filament network at CME sites may be less efficient at force production in the absence of Myo1E, as the distance between the center of the clathrin coat and the center of the Arp2/3 complex signal was greater in Myo1E knockout cells, possibly indicating less directed force being generated toward the invagination. These results suggest that Myo1E’s role in recruiting Arp2/3-activating NPFs is most crucial at high membrane tension, where CME sites need more Arp2/3 complex, and presumably more myosin motor activity to augment actin filament polymerization for efficient endocytosis. Consistent with Myo1E performing a local function at individual CME sites, we observed differences between sites that recruit Myo1E and those that recruit little or no Myo1E. Therefore, we aimed to identify conditions under which Myo1E becomes more prominent at CME sites. We found that in response to both mechanical and chemical perturbations that increase membrane tension, more Myo1E was recruited to CME sites. Moreover, Myo1E recruitment to CME sites increased in proportion to the magnitude of an osmotic shock. In addition, as the magnitude of the osmotic shock increased, CME sites with and without Myo1E tended to behave more similarly (both became longer lived and showed less movement in the plane of the plasma membrane) than sites in untreated cells. This observation is consistent with the possibility that, under increased membrane tension, all CME sites become longer lived and more stable, and that Myo1E is recruited to sites under the highest load to assist in vesicle formation. Additionally, we observed that CME sites that recruit Myo1E also tended to recruit more Dnm2, which showed longer lifetimes and increased fluorescent intensity. Together, these observations support the notion that sites that recruit Myo1E under normal physiological conditions are experiencing locally elevated membrane tension and might need additional force to achieve membrane invagination, with more Myo1E overall recruited as membrane tension is increased globally.

Much of the endocytic machinery is conserved between yeast and mammalian cells. Biochemical and biophysical analysis of the yeast type-1 myosin, Myo5, indicated that the motor activity of this myosin primarily generates power rather than forming tension-sensitive catch bonds, which some type-1 myosins form^42^. Using kinetic data from these experiments, recent simulations revealed that CME sites spend less time retracting and retract shorter distances when a motor with the parameters determined for Myo5 is present compared to a motor with stronger tension-sensitive catch bond behavior, such as mammalian Myo1b.^43–45^ Our observation of reduced MSD of Myo1E-positive endocytic sites in mammalian cells is consistent with these *in silico* findings. Due to the high turgor pressure in yeast, Myo5 is essential for all CME events. Our results suggest that, in mammalian cells, the class I myosin, Myo1E, plays a more significant role at sites of high membrane tension to maximize force generation for internalization. We hypothesize that, in the presence of Myo1E, more actin filaments polymerize against the plasma membrane, and assembly forces—possibly augmented by Myo1E’s motor activity—generate more directed and effective force to resist site retraction and promote the internalization of stalled endocytic sites.

## Acknowledgments

We thank Dr. Mira Krendel for sharing the Myosin1E antibody used for western blotting experiments. We thank Patrick Zager for helping build the micromanipulator device. We thank Robert Cail and Max Ferrin for their critical reading and feedback on this manuscript. This research was conducted with US Government support, under NIH Grant R35GM118149 to D.G.D and R35GM118167 O.D.W.

## Author Contributions

S.L.S and D.G.D designed research, S.L.S performed research and analyzed the data, T.Z. contributed reagents and performed 3D STORM imaging experiments, W.L. contributed analytical tools for 3D STORM co-localization data, H.DB. developed micro-aspirating assay and contributed analytical tools, Q.Z. helped collect live-cell imaging data for osmotic shock experiments. K.X. and O.W. provided guidance on experimental design and execution. S.L.S and D.G.D wrote the manuscript.

## Competing Interests

The authors declare no competing interest.

## Methods & Materials

### Cell Culture

The WTC10 hiPSC line was obtained from the laboratory of Bruce Conklin lab at UCSF. The hiPSCs were cultured on Matrigel (hESC-Qualified Matrix, Corning #354277) coated dishes in StemFlex medium (Thermo Fisher #A3349401) with Penicillin/Streptomycin at 37°C with 5% CO2. Cultures were passaged with Versene (Thermo Fisher #15040066) twice a week.

### Genome-editing

The MYO1E gene was edited in WTC10 hiPSCs in an AP2M1 dual-tagged TagRFP-T and DNM2 dual-tagged tagGFP2 background, which was previously edited ^4,20^. The Cas9-sgRNA complex electroporation method was used to edit the MYO1E gene to create in-frame fusions with the HaloTag. *S. pyogenes* NLS-Cas9 was purified in the University of California, Berkeley QB3 MacroLab. The MYO1E sgRNA (AACTATGTGACCAAGATCTG) was purchased from IDT. Sequence- and ligation-independent cloning (New England Biolabs) was used to construct the donor plasmid containing MYO1E 5’homology- gatccggtaccagcgatccaccggtcgccacc-HaloTag-MYO1E 3’homology. One week after electroporation (Lonza, Cat #: VOH-5012) of the Cas9-sgRNA complex and donor plasmid, the HaloTag positive cells were single cell sorted using a BD FACSAria Fusion Flow Cytometer (BD Biosciences) into Matrigel-coated 96-well plates with StemFlex medium with ROCK Inhibitor. Clones were confirmed by PCR and Sanger sequencing of the genomic DNA locus around the insertion site, as well as Western blot to check for protein expression. Both alleles of MYO1E were tagged with HaloTag in the hiPSC lines used.

The MYO1E gene was knocked out in the AP2- TagRFP-T; DNM2-tagGFP2 hiPSC line using the same technique as mentioned above. The ARPC3 gene was edited into this knockout cell line, also using Cas9. The MYO1E sgRNA (ACCAAGATCTGAGGTGCCCG) was purchased from Synthego. The ARPC3 sgRNA (CCTGGACAGTGAAGGGAGCC) was purchased from IDT. Sequence- and ligation-independent cloning (New England Biolabs) was used to construct the donor plasmid containing ARPC3 5’homology- ggatccggtaccagcgatccaccggtcgccacc-HaloTag-ARPC3 3’homology. Clones were confirmed by PCR and Sanger sequencing of the genomic DNA locus around the insertion site, as well as Western blot to check for protein expression. Both alleles of MYO1E were knocked out, and a single allele of ArpC3 was tagged with HaloTag in the hiPSC line used.

### Western Blots

For each sample, once cells in one well of a 6-well plate were 80-90% confluent, the plate was placed on ice and the media was removed, followed by a wash with cold 1X PBS. Cold lysis buffer (150 mM NaCl, 50 mM Tris pH 7.5, 1% NP-40, protease inhibitor (Roche, Cat#:11836170001)) was added to the well, and the cells were mechanically lifted off the plate with a cell scraper. Lysate was mixed with Laemmli buffer (0.5% BME)^46^. The samples were boiled at 95°C for 5 minutes and spun for 5 minutes at ∼17k x g. After cooling, the samples were loaded onto an SDS-PAGE gel and transferred to nitrocellulose membranes overnight for immunoblotting. Blots were blocked for 1 hour at room temperature in 5% dried milk in 1x TBST. Blots were then incubated overnight at 4°C in 0.5% TBST milk with primary antibodies targeting Myo1E (1:2,500, gift from Krendel lab), HaloTag (1:1000, Promega, Cat#: G9211), GAPDH (1:100,000 dilution, Proteintech, Cat#: 104941-AP). After primary antibody incubation, the blots were washed 3 times for 10 minutes in TBST. The blots were then incubated in the dark at room temperature for 1 hour with secondary antibodies in TBST (1:20,000 for both antibodies, Licor IRDye 800CW, Cat#:926-32212 or 926-32213)

### Total Internal Reflection Microscopy (TIRF) live-cell imaging

Two days before imaging, the hiPSCs were seeded onto Matrigel-coated cover glasses in 8-well chambers (Cellvis, Cat#: C8-1.5H-N). Two hours before imaging, the cells were incubated with 100 mM JF635-HaloTag ligand in StemFlex medium for 1 hour. After incubation, the cells were washed with StemFlex medium three times for 5 minutes at 37°C. The cells were imaged using a Nikon Ti2-E inverted microscope. During imaging, the cells were maintained at 37°C using a stage top incubator (Oko Labs) in DMEM/F12 + 10X StemFlex Supplement + 10 mM HEPES buffer. Images were acquired using a Nikon 60X CFI Apo TIRF objective (1.49 NA) and an Orca Fusion Genn III sCMOS camera (Hamamatsu) at 1x magnification using Nikon NIS Elements software. Using a LUNF 4-line laser launch (Nikon Instruments) and iLas2 TIRF/FRAP module (Gataca Systems), total internal reflection fluorescence (TIRF) illuminated a single focal plane of the field and was imaged every second for 5 minutes.

### Two-color 3D Stochastic Optical Reconstruction Microscopy (STORM) imaging/analysis STORM sample preparation

Cells were seeded onto 35-mm No. 1.5 poly-lysine coated MatTek dishes (MatTek, Cat#: P35GC-1.5-14-C) two days before fixation. Cells were fixed with 4% (vol/vol) paraformaldehyde (Electron Microscopy Sciences, Cat#: 15710) in Cytoskeleton Buffer (10 mM MES, 150 mM NaCl, 5 mM EGTA, 5 mM Glucose, 5 mM MgCl2, 0.005% NaN3, pH 6.1) at room temperature for 50 min and then washed twice for 5 minutes each with a freshly prepared 0.1% (wt/vol) NaBH4 solution in Dulbecco’s phosphate-buffered saline (DPBS) (Gibco, Cat#:14190144). Subsequently, the samples were washed 3 times for 10 min in DPBS. Samples were then blocked for 20 min in the blocking and permeabilization buffer [3% (wt/vol) BSA and 0.1% (wt/vol) saponin in DPBS]. The samples were then incubated with primary antibodies overnight at 4 °C: mouse anti-clathrin heavy chain (Invitrogen, MA1-065, 1:400 dilution) and rabbit anti-Halotag (Promega, G9281, 1:200 dilution). The next day, samples were washed three times in the washing buffer (0.1× blocking and permeabilization buffer in DPBS) for 10 min. Samples were then incubated with secondary antibodies in the blocking buffer for 60 min at room temperature in the dark, washed three times for 10 min in the washing buffer, and then three times for 10 min in DPBS. Secondary antibodies used were anti-rabbit labeled by Alexa Fluor 647 (AF647) (ThermoFisher, Cat#: A32787, 1:200 dilution) and anti-mouse (Jackson ImmunoResearch, 715-005-151) conjugated with amine-reactive succinimidyl ester CF597R (Biothium, 96092). Samples were washed three times with PBS before STORM imaging.

### Two-color 3D STORM imaging and analysis

STORM was performed as described previously on a homebuilt inverted microscope using a Nikon CFI Plan Apo λ 100x oil-immersion objective (NA = 1.45).^7,20,32,47^ The imaging buffer consisted of 5% (wt/vol) glucose, 100 mM cysteamine (Sigma-Aldrich, 30070), 0.8 mg/ml glucose oxidase (Sigma-Aldrich, G2133), and 40 μg/ml catalase (Sigma-Aldrich, C30) in 1 M Tris-HCl (pH 7.5)(Bates et al., 2007; Rust et al., 2006). For diffraction-limited epifluorescence images, lasers of 647 nm (for AF647), 560 nm (for CF597R), and 488 nm (for GFP) excited the sample at ∼10 W/cm^2^. For STORM imaging, lasers of 647 nm (for AF647) and 560 nm (for CF597R) excited the sample at ∼1.5 kW/cm^2^. ^32^ The angle of incidence was slightly below the critical angle of total internal reflection, thus illuminating a few micrometers into the sample. The relatively strong excitation powers switched most of the fluorescence molecules in the sample into a non-emitting dark state, leaving a low density of emitting single molecules in the view due to self-activation. A 405-nm laser was applied to further assist the activation of the dark-state molecules back to the emitting state, with gradually increasing power of 0-5 W/cm^2^ during the experiment, to sustain a suitable density of single molecules in the view. The resulting stochastic photoswitching of single-molecule fluorescence was recorded using an Andor iXon Ultra 897 EM-CCD camera at 110 frames per second. A total of ∼80,000 frames were recorded per image for each channel. For 3D localization, a cylindrical lens was inserted in the light path to create astigmatism for encoding the single-molecule depth (Z) information.^30^ The raw STORM data was processed using previously described methods to render super-resolution images.^30,31^ To quantify the distances between clathrin and Arpc3 or Myo1e signals, regions containing single CME sites associated with Arpc3 or Myo1E were cropped out, and the distance between the centroids of the two channels was calculated using MATLAB.

### Micro-aspiration Assay

Two days before imaging, the hiPSCs were seeded onto Matrigel-coated 35-mm No. 1.5 poly-lysine coated MatTek dishes (MatTek, Cat#: P35GC-1.5-14-C). The micropipette aspiration experiments were conducted using a micromanipulator (Narishige MM-188NE) mounted on the Nikon Ti2-E inverted microscope described above, with a custom 3D-printed mount. Two hours before imaging, the cells were incubated with 100 mM JF635-HaloTag ligand in StemFlex medium for 1 hour. After incubation, the cells were washed with StemFlex medium three times for 5 minutes at 37°C. While the cells were incubating with ligand, the micro-aspirating needle (Sunlight Medical, Cat#: SRP-05P-35) was coated with 5% BSA (wt/vol) in DMEM/F12 + 10X StemFlex Supplement for 10 minutes on the microscope stage to prevent cell adhesion. During imaging, the cells were maintained at 37°C with a stage top incubator (Oko Labs) in DMEM/F12 + 10X StemFlex Supplement + 10 mM HEPES buffer. Once a desired region of the hiPSC colony was identified, the needle was lowered to the imaging surface. The cells were imaged using the same setup as mentioned previously. After imaging had started, the cells were aspirated at a constant force using the CellTram 4r Air (Caliber Scientific) during imaging. For the analysis, the cells in the colony surrounding the cell being pulled were analyzed because the cell being aspirated was often lifted away from the TIRF field.

### Osmotic Shock Assay

Two days before imaging, the hiPSCs were seeded onto Matrigel-coated cover glasses in 8-well chambers (Cellvis, Cat#: C8-1.5H-N). Two hours before imaging, the cells were incubated with 100 mM JF635-HaloTag ligand in StemFlex medium for 1 hour. After incubation, the cells were washed with StemFlex medium three times for 5 minutes at 37°C. The cells were then put in isotonic imaging media (DMEM/F12 + 10X StemFlex Supplement + 10 mM HEPES buffer, 300 mOsm) at the desired volume to reach the ideal osmolarity with hypotonic solution (10 mM CaCl2 0.3 mM MgCl2 and 0.1 mM MgSO4). 10 mM HEPES buffer was present in all solutions. The 225 mOsm hypotonic imaging media was prepared by 1:3 vol/vol dilution with hypotonic solution to DMEM/F12 + 10X StemFlex Supplement. The 150 mOsm was made using a 1:1 vol/vol dilution, and the 75 mOsm was made using a 3:1 vol/vol dilution. The 300 mOsm control was added with 1:1 vol/vol of imaging media. One control movie was taken prior to the addition of hypotonic or control solution. Once the hypotonic or control solution was added to the desired well and gently mixed, imaging commenced 10 seconds later.

### Transferrin Uptake Assay

The hiPSCs were starved in HBSS + 0.5% BSA for 30 minutes at 37°C. After starvation, the cells were incubated with 10 µg/ml Tf–Alexa Fluor 647 at 37°C for the allotted amount of time in StemFlex. The cells were then transferred to ice, and the surface-bound Tf–Alexa Fluor 647 was removed using ice-cold acid buffer (HBSS, 0.5% acetic acid, and 0.5 M NaCl) for 45 seconds. The cells were then neutralized by gently washing with ice-cold PBS three times. The cells were harvested with Accutase (ThermoFisher Scientific, Cat#: A1110501) at 37°C for 2 minutes, resuspended in cold PBS, and spun at 4°C 300 rpm for 5 minutes. The supernatant was removed, and the cells were fixed in cold 4% PFA (vol/vol) in PBS at room temperature for 15 minutes. After fixation, the cells were spun at 4°C 300 rpm for 3 minutes, washed with cold PBS, spun again, and resuspended in cold PBS. The cells were put through a 35 µm cell strainer (ThermoFisher Scientific, Cat#: 08-771-23) before sorting. The cells were sorted for positive signal in the Allophycocyanin (APC), Fluorescein isothiocyanate (FITC), and Phycoerythrin (PE) channels using a BD Symphony A3 Flow Cytometer (BD Biosciences). The results were normalized to surface expression of the Tf receptor, which was evaluated by incubating the cells with Tf–Alexa Fluor 647 on ice for 30 min.

### TIRF image processing and analysis

Events, which are deemed as tracked diffraction-limited spots, were extracted using the MATLAB tracking package, cmeAnalysis and processed in Python Jupyter Notebooks ^29^. AP2-tagRFP-T was used as the fiducial marker for CME, and DNM2-tagGFP2 was used as a secondary channel to mark vesicle scission. The JF635 was tracked separately from paired AP2-RFP/Dnm2-GFP movies and linked to CCPs downstream of cmeAnalysis to allow for the determination of discrete nucleation and disassembly events of the protein in the third channel. The selection criteria for valid tracks were described previously in Jin et. al. 2022.

### Code Availability & Image Analysis tools

https://github.com/DrubinBarnes

---

**Figure S1.**
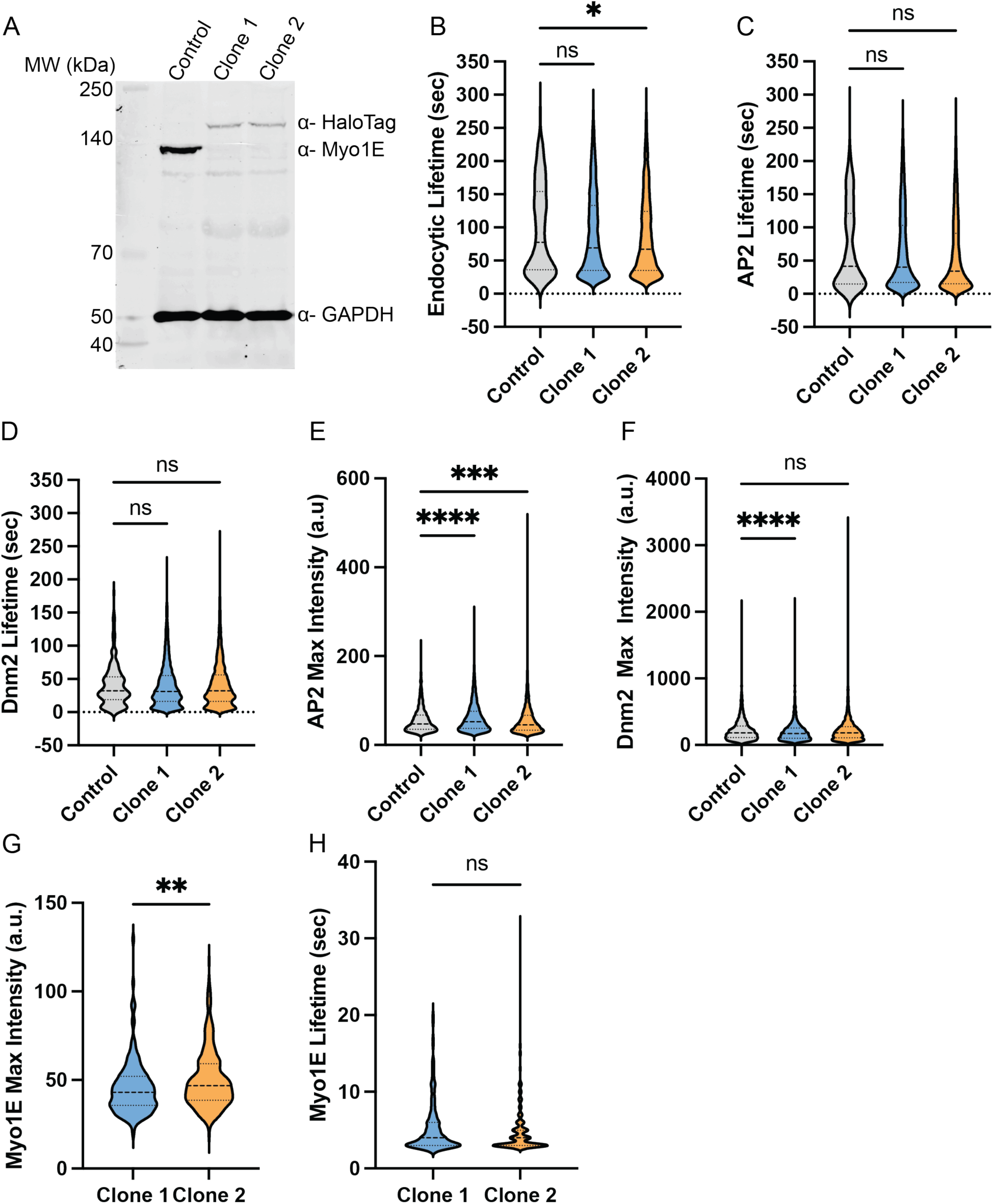
(A) Western blot of Myo1E-HaloTag knock-in. AP2-tagRFP-T; DNM2-tagGFP2 hiPSC line is the parent control cell line. Myo1E-HaloTag is 160 kDa, Myo1E endogenous is 127 kDa and GAPDH is 36 kDa. (B-G) Volin plots comparing control AP2-tagRFP-T; DNM2-tagGFP2 hiPSCs with two AP2-tagRFP-T; DNM2-tagGFP2; MYO1E-JF635 clones generated from clonal expansion after genome-editing MYO1E with a HaloTag. Clone 2 was used for these studies as it had a more similar Dnm2-GFP maximum intensity to the control compared to Clone 1.

**Figure S2.**
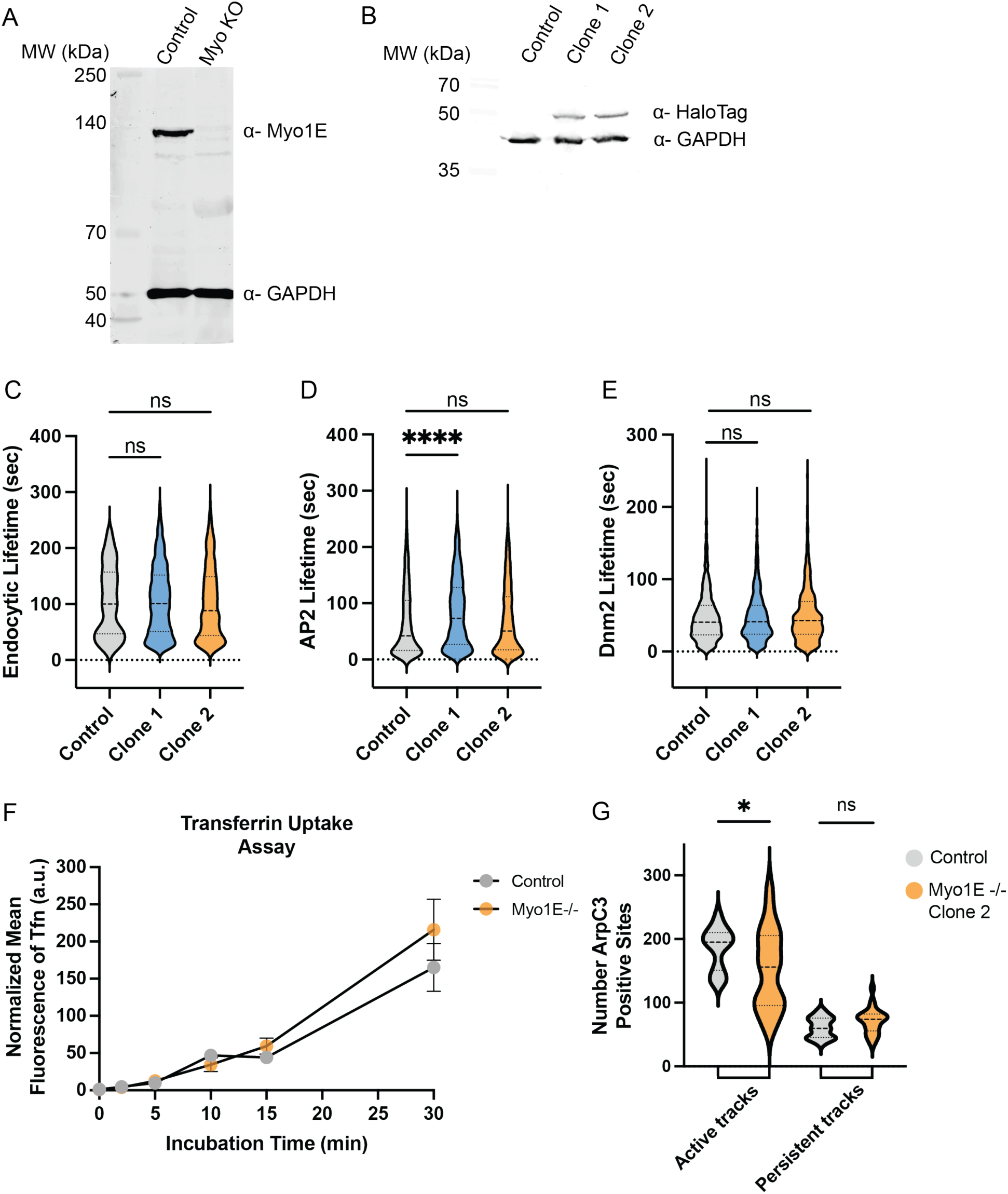
(A) Western blot of AP2-tagRFP-T; DNM2-tagGFP2 control hiPSCs and homozygous Myo1E knockout hiPSCs in an AP2-tagRFP-T; DNM2-tagGFP2 background. Myo1E endogenous is 127 kDa and GAPDH is 36 kDa. (B) Western blot of AP2-tagRFP-T; DNM2-tagGFP2 control hiPSCs and ArpC3-HaloTag knock-in in Myo1E knockout hiPSCs. ArpC3-HaloTag is 54 kDa and GAPDH is 36 kDa. (C-E) Volin plots comparing control AP2-tagRFP-T; DNM2-tagGFP2; ArpC3-HaloTag hiPSCs with two AP2-tagRFP-T; DNM2-tagGFP2; ArpC3-HaloTag; MYO1E knockout clones generated from clonal expansion after knocking out MYO1E and tagging ArpC3 with a HaloTag. Clone 2 was used for these studies as it had more similar AP2-RFP lifetimes compared to the control. (F) Time-course transferrin-647 uptake assay of control cells compared to knockout cells. (G) Violin plot showing the number of endocytic sites in control cells and Myo1E-/- cells for both active sites and persistent sites.

**Figure S3.**
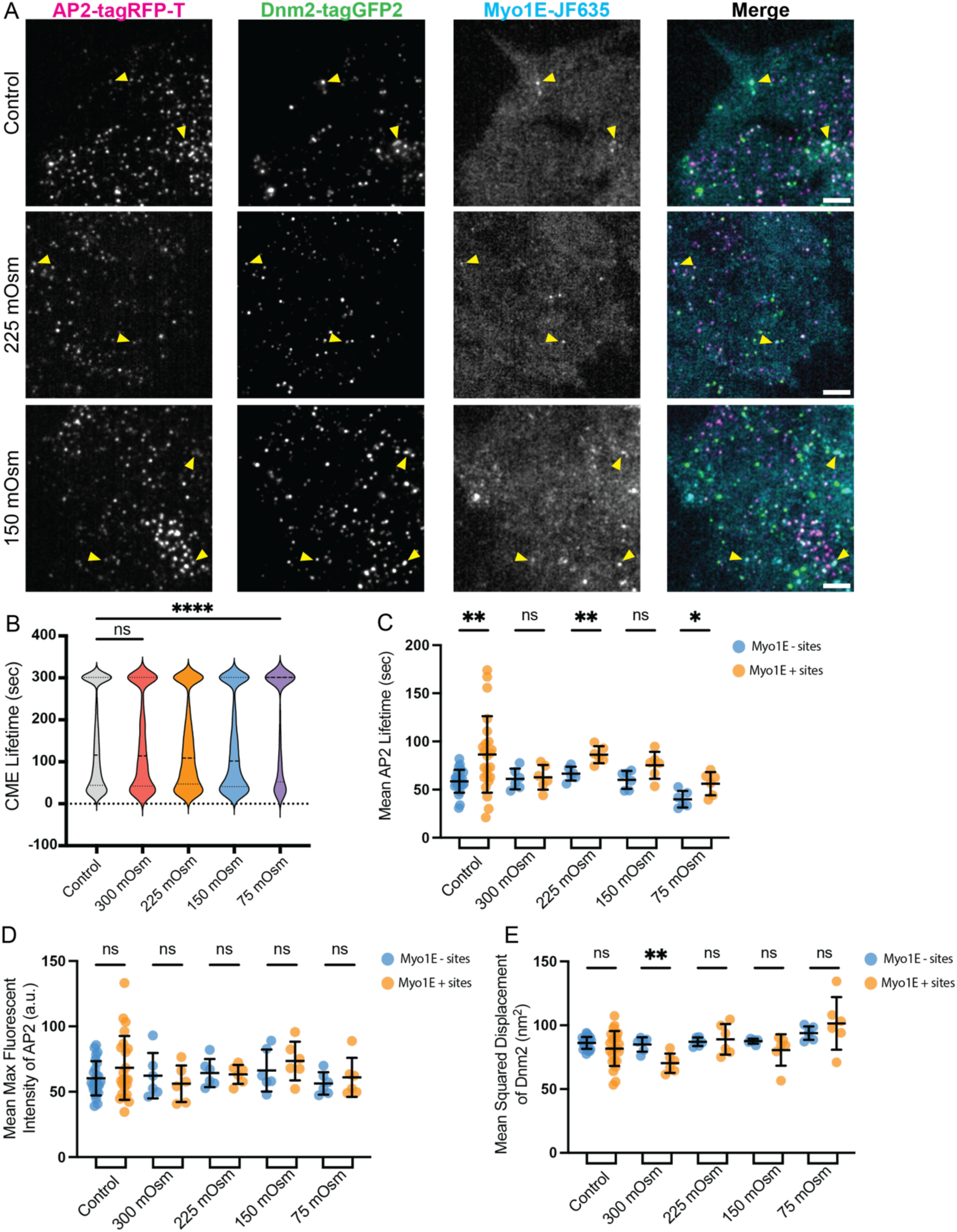
(A) TIRF image of ADM cells after the addition of hypotonic media to reach a final concentration of 225 and 150 mOsm. Scale bar = 5 µm. (B) Violin plot of CME lifetimes of all AP2-RFP;DNM2-GFP sites under varying osmotic shock conditions (P****< 0.0001, for Kruskal-Wallis). (C-D) Scatterplots characterizing AP2-RFP lifetime, mean fluorescent intensity, and mean displacement between endocytic sites that are Myo1E negative and Myo1E positive. (P**** < 0.0001, for Mann-Whitney). Error bars show SD.

**Movie 1.** TIRF movie of hiPSCs endogenously expressing AP2-tagRFP-T (magenta), DNM2-tagGFP2 (green), and Myo1E-JF635 (cyan). Related to Fig. 1 (A-D). Frame rate is 1 frame per second (fps) and played back at 10 fps. Scale bar = 5 µm.

**Movie 2.** TIRF movie of hiPSCs endogenously expressing AP2-tagRFP-T (magenta), DNM2-tagGFP2 (green), and ArpC3-JF635 (cyan). Related to Fig. 2 (A-E). Cells are in a wildtype background. Frame rate is 1 frame per second (fps) and played back at 10 fps. Scale bar = 5 µm.

**Movie 3.** TIRF movie of hiPSCs endogenously expressing AP2-tagRFP-T (magenta), DNM2-tagGFP2 (green), and ArpC3-JF635 (cyan). Related to Fig. 2 (A-E). Cells are in a Myo1E-/- background. Frame rate is 1 frame per second (fps) and played back at 10 fps. Scale bar = 5 µm.

**Movie 4.** TIRF movie of hiPSCs endogenously expressing AP2-tagRFP-T (magenta), DNM2-tagGFP2 (green), and ArpC3-JF635 (cyan). Related to Fig. 3 (A-C). Cells are in a wildtype background. Frame rate is 1 frame per second (fps) and played back at 10 fps. Scale bar = 5 µm.

**Movie 5.** TIRF movie of hiPSCs endogenously expressing AP2-tagRFP-T (magenta), DNM2-tagGFP2 (green), and Myo1E-JF635 (cyan). Related to Fig. 3 (D-F). Cells are in a wildtype background. Frame rate is 1 frame per second (fps) and played back at 10 fps. Scale bar = 5 µm.

**Movie 6.** TIRF movie of hiPSCs endogenously expressing AP2-tagRFP-T (magenta), DNM2-tagGFP2 (green), and Myo1E-JF635 (cyan) in 300 mOsm. Related to Fig. 4 (A-G). Cells are in a wildtype background. Frame rate is 1 frame per second (fps) and played back at 10 fps. Scale bar = 5 µm.

**Movie 7.** TIRF movie of hiPSCs endogenously expressing AP2-tagRFP-T (magenta), DNM2-tagGFP2 (green), and Myo1E-JF635 (cyan) in 75 mOsm. Related to Fig. 4 (A-G). Cells are in a wildtype background. Frame rate is 1 frame per second (fps) and played back at 10 fps. Scale bar = 5 µm.

**Movie 8.** TIRF movie of hiPSCs endogenously expressing AP2-tagRFP-T (magenta), DNM2-tagGFP2 (green), and ArpC3-JF635 (cyan) in 300 mOsm. Related to Fig. 5 (A, C-G). Cells are in a wildtype background. Frame rate is 1 frame per second (fps) and played back at 10 fps. Scale bar = 5 µm.

**Movie 9.** TIRF movie of hiPSCs endogenously expressing AP2-tagRFP-T (magenta), DNM2-tagGFP2 (green), and ArpC3-JF635 (cyan) in 75 mOsm. Related to Fig. 5(A, C-G). Cells are in a wildtype background. Frame rate is 1 frame per second (fps) and played back at 10 fps. Scale bar = 5 µm.

**Movie 10.** TIRF movie of hiPSCs endogenously expressing AP2-tagRFP-T (magenta), DNM2-tagGFP2 (green), and ArpC3-JF635 (cyan) in 300 mOsm. Related to Fig. 5 (B, C-G). Cells are in a Myo1E-/- background. Frame rate is 1 frame per second (fps) and played back at 10 fps. Scale bar = 5 µm.

**Movie 11.** TIRF movie of hiPSCs endogenously expressing AP2-tagRFP-T (magenta), DNM2-tagGFP2 (green), and ArpC3-JF635 (cyan) in 75 mOsm. Related to Fig. 5 (B, C-G). Cells are in a Myo1E-/- background. Frame rate is 1 frame per second (fps) and played back at 10 fps. Scale bar = 5 µm.

## Notes

### Competing Interest Statement

The authors have declared no competing interest.

