## Supplementary figures and images for "Class-I myosin responds to changes in membrane tension during clathrin-mediated endocytosis in human induced pluripotent stem cells"

### Figure S1

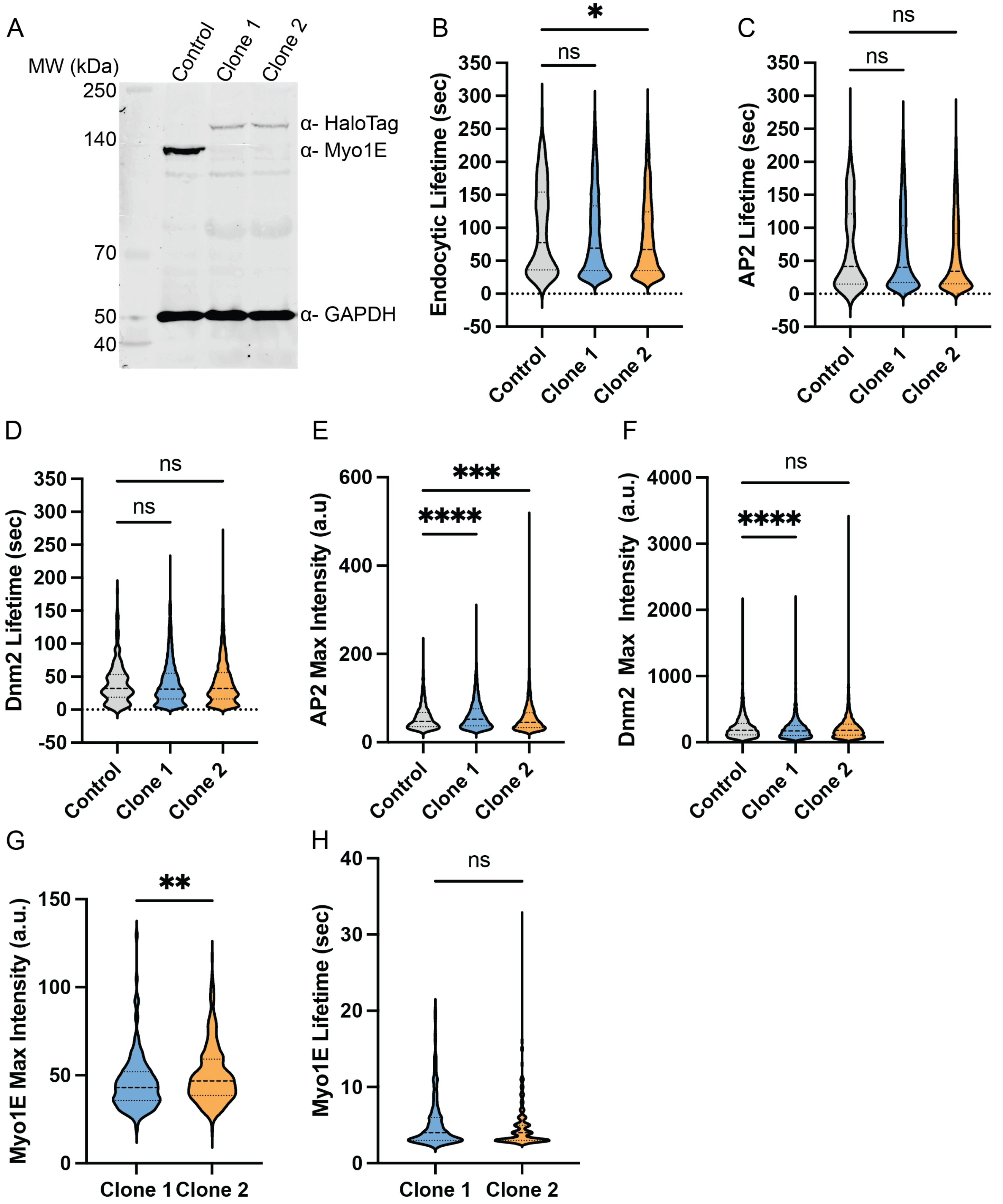

### Figure S2

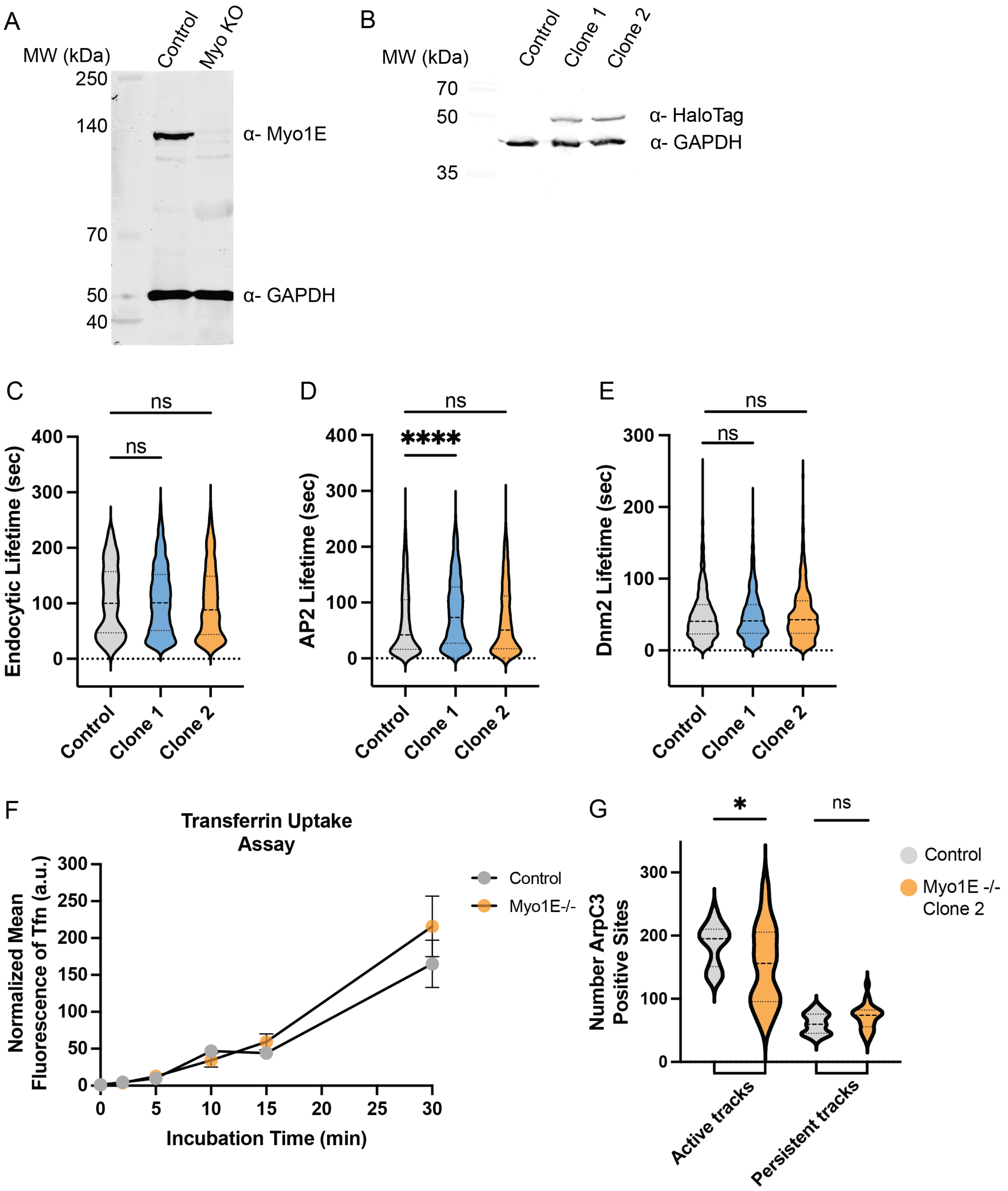

### Figure S3

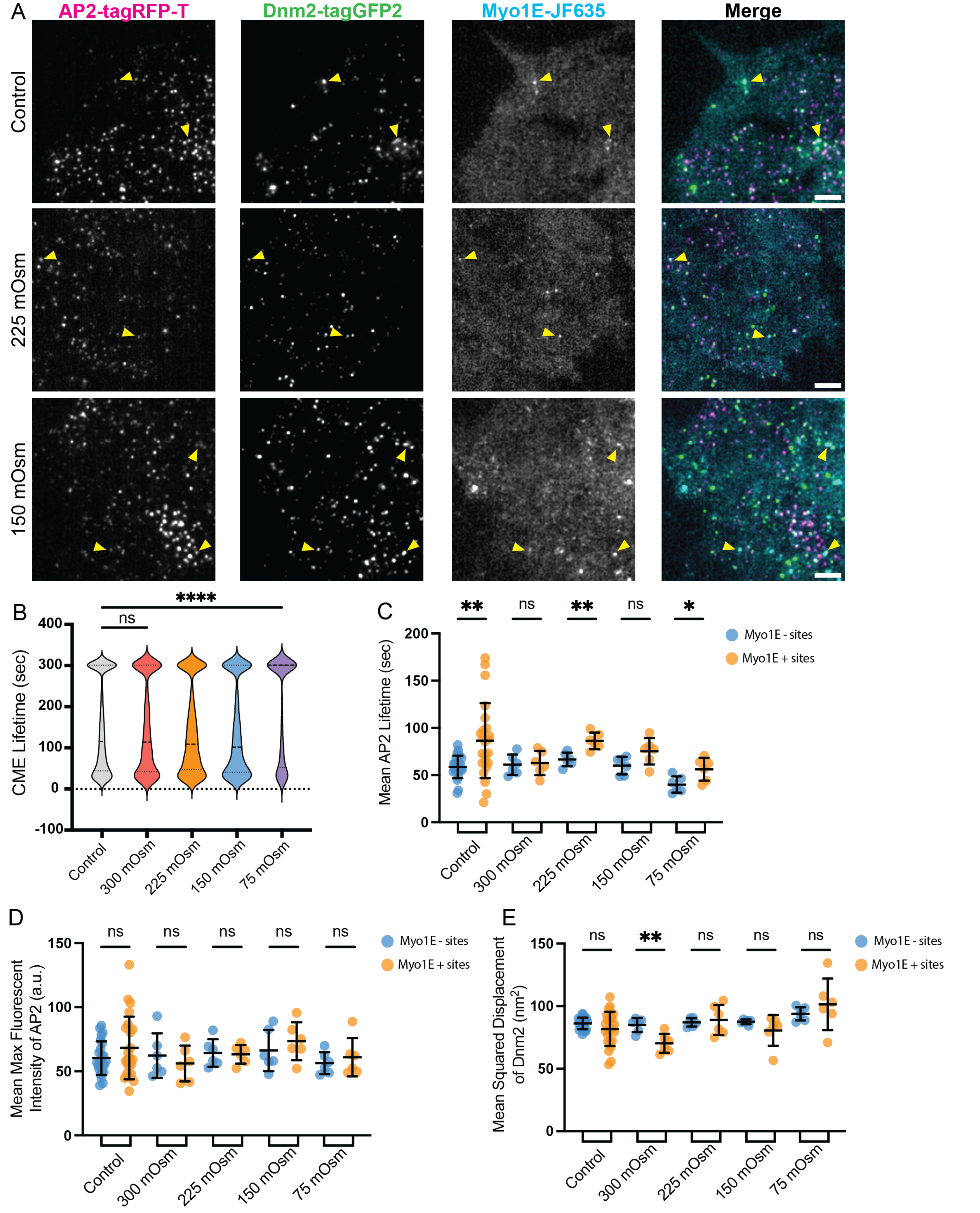
